# The marbled crayfish as a paradigm for saltational speciation by autopolyploidy and parthenogenesis in animals

**DOI:** 10.1101/025254

**Authors:** Günter Vogt, Cassandra Falckenhayn, Anne Schrimpf, Katharina Schmid, Katharina Hanna, Jörn Panteleit, Mark Helm, Ralf Schulz, Frank Lyko

**Author notes:** Authors for correspondence: Günter Vogt Frank Lyko. present address: Faculty of Biosciences, University of Heidelberg, Im Neuenheimer Feld 230, 69120 Heidelberg, Germany (Günter Vogt).

## Abstract

The parthenogenetic all-female marbled crayfish is a novel research model and potent invader of freshwater ecosystems. It is a triploid descendant of the sexually reproducing slough crayfish, *Procambarus fallax*, but its taxonomic status has remained unsettled. By cross-breeding experiments and parentage analysis we show here that marbled crayfish and *P. fallax* are reproductively separated. Both crayfish copulate readily, suggesting that the reproductive barrier is set at the cytogenetic rather than the behavioural level. Analysis of complete mitochondrial genomes of marbled crayfish from laboratory lineages and wild populations demonstrates genetic identity and indicates a single origin. Flow cytometric comparison of DNA contents of haemocytes and analysis of nuclear microsatellite loci confirm triploidy and suggest autopolyploidization as its cause. Global DNA methylation is significantly reduced in marbled crayfish implying the involvement of molecular epigenetic mechanisms in its origination. Morphologically, both crayfish are very similar but growth and fecundity are considerably larger in marbled crayfish, making it a different animal with superior fitness. These data and the high probability of a divergent future evolution of the marbled crayfish and *P. fallax* clusters suggest that marbled crayfish should be considered as an independent asexual species. Our findings also establish the *P. fallax-*marbled crayfish pair as a novel paradigm for rare chromosomal speciation by autopolyploidy and parthenogenesis in animals and for saltational evolution in general.

## 1. Introduction

In the last decade, the marbled crayfish (Marmorkrebs) has gained considerable attention in the scientific community and the public because of its obligatory parthenogenetic reproduction, its suitability as a research model and its high potential as an invasive species [1-9]. It was discovered in 1995 in the German aquarium trade [2] and has become a popular pet in Europe and other continents since then [10,11]. Thriving wild populations have meanwhile developed from releases in several European countries and Madagascar and are feared to threaten native crayfish species by competition and transmission of the crayfish plague [7-9,12,13].

By comparison of morphological traits and molecular markers, Martin and colleagues [14] have identified the sexually reproducing slough crayfish *Procambarus fallax* from Florida and southernmost Georgia as the mother species of marbled crayfish. However, its taxonomic position remained unsettled. Martin *et al.* [14] suggested the provisional name *Procambarus fallax* forma *virginalis*, being aware that forma is not a valid category in animal taxonomy. Meanwhile, several important characteristics of marbled crayfish have been described in detail, including morphology [12], embryonic development [15,16], life history [16-19], parthenogenetic reproduction [1,20,21] and a triploid karyotype [22].

Speciation in parthenogenetic lineages is a problematic issue because parthenogens do not fit into the classical concepts of speciation, as discussed in detail by Mayr [23], Coyne and Orr [24], Barraclough *et al.* [25], Birky and Barraclough [26] and Martin *et al.* [14]. However, Barraclough and colleagues emphasized the importance of understanding diversification and speciation in asexual organisms, not least to test theories about the evolutionary advantage of sex [25,26]. They provided a theory on speciation in asexuals, which they named Evolutionary Genetic Species Concept [26]. This theory focuses on the criterion that the individuals of the parent species and the neo-species form discrete clusters of very similar genotypes and phenotypes. The new cluster should be of a single origin and both clusters must be separated from each other by reproductive or geographic isolation and a gap of genetic and phenotypic traits so that natural selection can ensure a divergent evolution over time [25-28].

Stimulated by the paper by Martin *et al.* [14] there is an ongoing discussion among marbled crayfish experts whether this animal should be treated as a parthenogenetic lineage of *P. fallax* or a species in its own right. In order to examine this issue in detail we have tested the above listed operational definitions for asexual species with several experimental and technical approaches. Cross-breeding experiments between marbled crayfish and slough crayfish and parentage analysis by microsatellite markers were performed to test for reproductive isolation. Complete mitochondrial genomes and nuclear microsatellite patterns of marbled crayfish from several laboratory lineages and wild populations were analysed to clarify single origin and to establish its genotypic characteristics. The DNA content of haemocytes, mitochondrial genome sequences and microsatellite patterns was compared between marbled crayfish, *P. fallax* and the closely related *Procambarus alleni* to obtain information about the mode of triploidization of the marbled crayfish. Global DNA methylation was determined to examine the involvement of epigenetic mechanisms in speciation. Finally, taxonomically relevant morphological characters and ecologically and evolutionarily important life history traits were compared to reveal phenotypic differences between the marbled crayfish and *P. fallax* clusters.

## 2. Material and methods

### 2.1 Animals

The following animals were used: (1) marbled crayfish *Procambarus fallax* (Hagen, 1870) f. *virginalis* from our laboratory lineages named “Heidelberg” and “Petshop” and from two wild populations in Germany and Madagascar, (2) *Procambarus fallax* (Hagen, 1870) from our laboratory population and the aquarium trade, (3) *Procambarus alleni* (Faxon, 1884) from the aquarium trade, and 4) *Procambarus clarkii* (Girard, 1852) from an invasive Swiss population. The Heidelberg lineage was founded by G.V. in February 2003 from a single female, which originated from the oldest documented marbled crayfish aquarium population founded in 1995 by F. Steuerwald. The Petshop lineage was established by G.V. in February 2004 from a single female purchased in a pet shop. The wild marbled crayfish were from Lake Moosweiher, Germany (provided by M. Pfeiffer), and a market in Antananarivo, Madagascar (provided by F. Glaw). Our *P. fallax* laboratory population was founded in February 2014 by a single pair obtained from the aquarium trade. All crayfish were raised under the same conditions. Animals were kept either individually or communally in plastic containers of 30×25×20 cm equipped with gravel and shelters. Tap water was used as the water source and replaced once a week. Water temperature was maintained at 20°C. All animals were fed with TetraWafer Mix pellets.

### 2.2 Cross-breeding experiments

For the 38 crossbreeding experiments we used three *P. fallax* males with total lengths (TL=tip of rostrum to end of telson) of 3.1-5.2 cm, five *P. fallax* females with TLs of 3.5-4.2 cm, 14 marbled crayfish females with TLs of 4.0-6.3 cm and two *P. alleni* males with TLs of 5.1-5.3 cm. All males were in the reproductively competent Form I as indicated by the presence of hooks on the ischia of the 3rd and 4th peraeopods. Eight of the 14 marbled crayfish females and 4 of the 5 *P. fallax* females had well-developed glair glands on the underside of the pleon indicating ovarian maturity and receptiveness. The behavioural experiments were performed in aquaria with an area of 26x16 cm without shelter. Pairs were observed for 2 hours and copulation was regarded as successful when the partners remained in typical copulation position for more than 10 min. Parentage of the offspring was determined by microsatellite analysis.

### 2.3 Microsatellite analysis

For microsatellite analysis, walking legs of specimens were fixed in 80% ethanol prior to extraction of nuclear DNA with the Blood& Cell Culture DNA Kit (Genomic Tips) from Qiagen (Hilden, Germany). A total of five microsatellite primer pairs were tested. Four of them were originally designed for *P. clarkii* (PclG-02, PclG-04, PclG-08, PclG-48) [29] and one pair (PclG-26) was designed for marbled crayfish based on the *P. clarkii* sequences [21]. The same microsatellite loci were additionally investigated in *P. alleni* and *P. clarkii*. PCR was carried out using a Primus 96 Cycler (Peqlab Biotechnologie, Erlangen, Germany). Fragment analysis was performed on a Beckman Coulter CEQ 8000 eight capillary sequencer (Beckman Coulter, Krefeld, Germany) using the Beckman Coulter DNA Size Standard Kit 400 bp. Loci were scored with GeneMarker, v.2.6 (SoftGenetics, State College, Pennsylvania, USA).

### 2.4 Sequencing, assembly and comparison of mitochondrial genomes

For comparison of complete mitochondrial genomes we used two cultured marbled crayfish from the Heidelberg and Petshop lineages, two wild marbled crayfish from Lake Moosweiher and Madagascar, one *P. fallax* female and one *P. alleni* female. DNA was isolated from hepatopancreases and abdominal musculature as described above and sequenced on an Illumina HiSeq platform. Read pairs were quality trimmed (quality value ≥30, minimum length ≥30) and the mitochondrial genome of the Heidelberg animal was assembled by Velvet2.0 [30]. The sequences of the other specimens were established by mapping against the Heidelberg sequence using Bowtie2 [31]. For the identification of single nucleotide polymorphisms (SNPs) between the marbled crayfish populations, we used mpileup and bcftools from SAMtools [32], requiring a quality value >30 for SNP calling. Mitochondrial genome sequences of *P. fallax* and *P. alleni* were generated by MITObim1.6 [33] using published mitochondrial DNA fragments from *P. fallax* (FJ619800) and *P. alleni* (HQ171462, FJ619802, HQ171451) as seed sequences. Mismatches in comparison to marbled crayfish sequences were identified by blastn alignments.

### 2.5 Measurement of DNA content by flow cytometry

Flow cytometry was used to determine the DNA content in haemocytes of *P. fallax* and marbled crayfish. Haemolymph was withdrawn through the articulating membrane between coxa and basis of the chelipeds, mixed 1:1 with crayfish anticoagulant buffer solution (100 mM glucose, 34 mM trisodium citrate, 26 mM citric acid, 15.8 mM EDTA, pH 4.6) and centrifuged for 5 min at 1400 rpm. The pellet was washed and re-suspended with 100 μl PBS. Samples were either stored in 10% DMSO at -80°C or immediately used for analysis of the DNA content. For flow cytometry 4 μl RNase A (Sigma-Aldrich, Munich, Germany) stock solution (50 mg/ml) was added to the samples and incubated for 5 min at room temperature followed by an incubation for 60 min with 5 μl propidium iodide (Life Technologies, Darmstadt, Germany) stock solution (1 mg/ml). The samples were then mixed 1:1 with PBS and the DNA-related fluorescence intensities of single cells were measured on a BD Accuri C6 Cytometer (BD Sciences, Heidelberg, Germany) with blue laser 488 nm and detection filter FL2 585/40 nm.

### 2.6 Measurement of global DNA methylation by mass spectrometry

Global DNA methylation was determined in three whole juveniles and selected tissues (hepatopancreas, abdominal musculature and ovary) of three adults of marbled crayfish and *P. fallax*. Sample preparation and LC-MS/MS analyses were conducted as previously described [34] and were performed on an Agilent 1260 LC system connected to an Agilent 6460 TripleQuad masspectrometer (Agilent, Böblingen, Germany). Briefly, after enzymatic hydrolysis to nucleosides, the samples were spiked with 250 fmol [D_3_]-5-methylcytosine as internal standard. The mass transitions resulting from the loss of desoxyribose (5-methylcytidine: 242 Th **→** 126 Th, [D_3_]-5-methylcytidine: 245 Th **→** 129 Th) by collision induced dissociation (CID) were analysed in dynamic multiple reaction monitoring mode (DMRM). Calibration curves using a stable isotope labelled internal standard were established for quantification of 5-methylcytidine. The linear regressions resulting from the double logarithmic plots were used to correlate the respective signals from LC-MS/MS analysis to known amounts of substance. The yield of detected modification was normalized to guanosine content (as equivalent to cytidine content) because of better signal quality. To assess the amount of guanosine, the areas of the DAD results, gained during the LC analysis, were correlated to their respective amounts of substance in the same way as above.

### 2.7 Investigation of morphological characters and life history traits

For comparison of morphological characters between marbled crayfish and *P. fallax* we used marbled crayfish with TLs of 4.0-8.4 cm and body weights of 1.4-15.2 g and *P. fallax* females with TLs of 3.6-5.7 cm and weights of 1.1-4.5 g. We focussed on annulus ventralis (sperm receptacle), areola of the carapace, cheliped chelae and coloration, the taxonomically most relevant characters in female Cambaridae [35-37]. For comparison of life history traits we analysed growth, time of sexual maturity, body size and clutch size. Growth was determined in batches raised under the same conditions by measurement of carapace length (CL), total length (TL) and body weight. Sexual maturity was deduced from the presence of glair glands. Mean and maximum body and clutch sizes were taken from our laboratory animals and published data on wild marbled crayfish and *P. fallax*.

## 3. Results

### 3.1 Crossbreeding experiments and parentage analysis

Crossbreeding experiments were performed to investigate whether marbled crayfish and *P. fallax* can interbreed and produce viable offspring. Behavioural observations revealed that marbled crayfish females and *P. fallax* males recognize each other as sexual partners. Courtship and mating behaviour included frontal approach, tearing with the chelipeds, intense sweeping with the antennae, sudden turning of the female and mounting by the male (figure 1). This courtship behaviour is also typical of other *Procambarus* species [38]. *P. fallax* males copulated with marbled crayfish females in 15 of 21 trials (71%) and with *P. fallax* females in 6 of 8 trials (86%) (table 1). In the marbled crayfish *x P. fallax* pairs, the first contact was often initiated by the marbled crayfish females. Some matings lasted for more than 1 hour. *P. fallax* males can turn significantly larger marbled crayfish females on the back but are not long enough to simultaneously fix the female’s chelipeds and insert the gonopods into the annulus ventralis. *P. alleni* males copulated neither with *P. fallax* nor with marbled crayfish females (table 1) suggesting that they did not recognize them as sexual partners.

**Figure 1.**
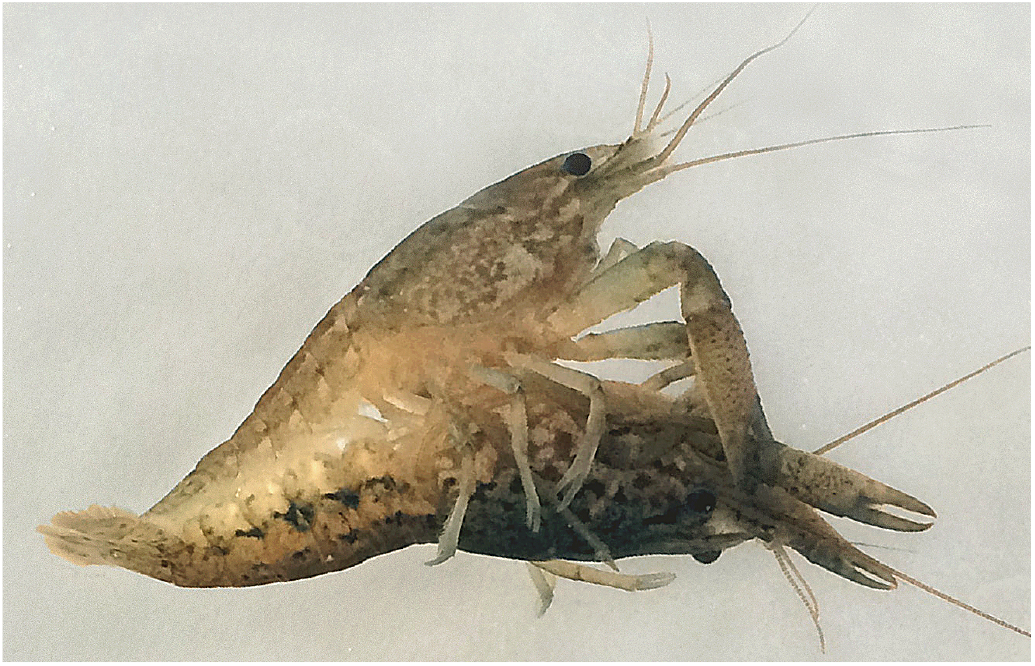
Mating of marbled crayfish female with *P. fallax* male. The male (top) holds the female firmly with the chelipeds and ischial hooks and his gonopods are plugged into the female’s spermatheca.

**Table 1.** Crossbreeding experiments between marbled crayfish, *P. fallax* and *P. alleni*.

| Males | Marbled crayfish females |  |  |  |  |  |  |  |  |  |  |  |  |  | *P. fallax* females |  |  |  |  |
| --- | --- | --- | --- | --- | --- | --- | --- | --- | --- | --- | --- | --- | --- | --- | --- | --- | --- | --- | --- |
|  | 1 | 2 | 3 | 4 | 5 | 6 | 7 | 8 | 9 | 10 | 11 | 12 | 13 | 14 | P1 | P2 | P3 | P4 | P5 |
| *P. fallax* 1 | x | x | x | x | xx |  | x | xo | o | x |  | x |  | o | x | x |  |  | x |
| *P. fallax* 2 |  |  |  |  |  | o | x |  | o | x | o | x |  |  |  |  | o | x |  |
| *P. fallax* 3 |  |  |  |  |  |  |  | x |  |  |  |  | x |  |  | x |  |  | x |
| *P. allenii* 1 |  |  |  |  | o |  | o | oo |  |  |  |  |  |  |  |  |  |  | o |
| *P. allenii* 2 |  |  |  |  |  |  | oo | oo |  |  |  |  |  |  |  |  |  |  | o |

x: mating; o: no mating; two letters: results of two trials.

We obtained a total of ten clutches from the crossbreeding experiments, eight from crosses of three *P. fallax* males with eight marbled crayfish females and two from crosses of two *P. fallax* males with two *P. fallax* females. Four of the *P. fallax x* marbled crayfish clutches and one *P. fallax x P. fallax* clutch developed into juveniles whereas the others decayed during embryonic development. In the *P. fallax x P. fallax* clutch we counted 10 females and 9 males at juvenile stage 7, reflecting the typical 1:1 sex ratio of sexually reproducing crayfish [39]. In contrast, in the four marbled crayfish *x P. fallax* batches the 6, 12, 61 and 93 analysed stage 7 offspring were all females indicating reproduction by parthenogenesis.

The progeny of our crossbreeding experiments were also investigated by microsatellite analysis to further clarify parentage. Microsatellite analysis is an established approach to assess parentage and geographic structuring in crayfish populations and to identify clonal lineages, triploids and hybrids [40-43]. Of the five primer pairs tested, three revealed PCR products that could be used for fragment length determination in marbled crayfish and *P. fallax*, namely PclG-02, PclG-04 and PclG-26. PclG-02 and PclG-26 were polymorphic and thus suitable for parentage testing. The microsatellite allele combinations in the analysed family groups of marbled crayfish females 1-4 x *P. fallax* male 1 were identical between mothers and offspring, namely 267 bp/271 bp/303 bp at locus PclG-02 and 189 bp/191 bp at locus PclG-26, but differed from the allele combination of the male that was 255 bp/267 bp and 185 bp/207 bp, respectively (table 2). All measurements were repeated at least twice, and in the case of the unusual PclG-02 up to five times per specimen. Our data indicate that the male did not contribute to the genome of the offspring and that the progeny is the product of apomictic parthenogenesis. The microsatellite patterns were not only identical between mother and offspring but also between the four batches (table 2) demonstrating clonality of all marbled crayfish from our laboratory.

**Table 2.** Parentage analysis in crossbreeds of marbled crayfish *x P. fallax*.

| Specimens | Microsatellite loci |  |
| --- | --- | --- |
|  | PclG-02 | PclG-26 |
| *P. fallax* father 1 | 255/267 | 185/207 |
| Marbled crayfish mothers 1-4 | 267/271/303 | 189/191 |
| Offspring of mother 1 (n=6) | 267/271/303 | 189/191 |
| Offspring of mother 2 (n=5) | 267/271/303 | 189/191 |
| Offspring of mother 3 (n=6) | 267/271/303 | 189/191 |
| Offspring of mother 4 (n=3) | 267/271/303 | 189/191 |

Values indicate fragment lengths in base pairs.

The *P. fallax* male 1 *x P. fallax* female 1 family was used as a positive control. Analysis of locus PclG-26 revealed the allele combinations 185 bp/207 bp in the father, 179 bp/185 bp in the mother and 179 bp/185 bp (2x), 179 bp/207 bp (4x), 185 bp/185 bp (4x) and 185 bp/207 bp (4x) in the 14 offspring. These data indicate Mendelian distribution and demonstrate that both parents contributed equally to the genome of the offspring, as is expected for sexually reproducing species.

### 3.2 Single origin and clonality of marbled crayfish populations

For a more detailed genetic analysis of marbled crayfish, we established complete mitochondrial genome sequences of specimens from our Heidelberg and Petshop lineages and from wild populations of Lake Moosweiher (Germany) and Madagascar by high-coverage shotgun sequencing and sequence mapping. Remarkably, these mitochondrial genome sequences were completely identical (figure 2), thus confirming the clonal nature of the tested populations and their single origin. Comparison of our sequences with the mitochondrial genome sequence of marbled crayfish published earlier by Shen *et al.* [44] revealed 6 scattered mismatches and major differences in one fragment ranging from position 4600 to 5500. These differences are probably related to technical issues because Shen and colleagues used PCR-based methods and primer walking single/double strands sequencing [44] whereas we used next-generation sequencing with a sequencing coverage per nucleotide of >100x.

**Figure 2.**
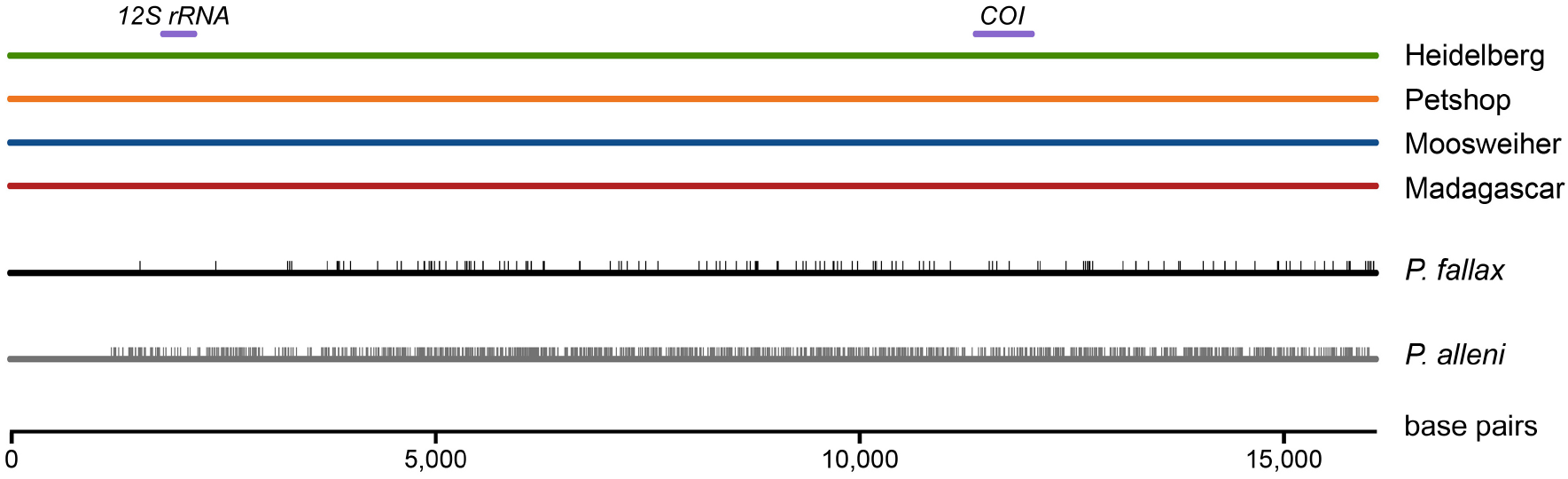
Comparison of complete mitochondrial genomes of marbled crayfish, *P. fallax* and *P. alleni*. The sequences of marbled crayfish from two laboratory populations (Heidelberg, Petshop) and two wild populations (Moosweiher, Madagascar) are completely identical. In contrast, the sequences of *P. fallax* and *P. alleni* differ in 144 and 1165 SNPs (vertical lines) from marbled crayfish, respectively. Purple bars indicate positions of *12S rRNA* and *cytochrome oxidase subunit I* (COI) genes that were earlier used for phylogenetic analysis [14].

We also established complete mitochondrial genome sequences for *P. fallax* and *P. alleni*. Analysis of the mitochondrial *12S rRNA*, *16S rRNA* and *cytochrome oxidase subunit I* genes have earlier indicated a close relationship between marbled crayfish and these species [1,7,14]. *P. alleni* occurs sympatrically with *P. fallax* in many locations in Florida [45] and was therefore regarded as a candidate that might have contributed to the origination of marbled crayfish by hybridization with *P. fallax* [46]. Sequence comparison revealed 144 single nucleotide polymorphisms (SNPs) between marbled crayfish and *P. fallax* but 1165 SNPs between marbled crayfish and *P. alleni* (figure 2). Interestingly, these SNPs were not evenly distributed over the mitochondrial genome, which explains why in the study by Martin *et al.* [14] small genetic differences between marbled crayfish and *P. fallax* were detected in the *cytochrome oxidase subunit I* gene but not in the *12S rRNA* gene. Our results confirm the close genetic relationship between marbled crayfish and *P. fallax* and a greater distance towards *P. alleni*.

The single origin and clonality of marbled crayfish from the laboratory and the wild was further confirmed by the analysis of microsatellite loci PclG-02, PclG-04 and PclG-26 in 24 specimens from our laboratory lineages (see parentage analysis), six specimens from a stable wild population in Lake Moosweiher [47] and one specimen from Madagascar [7]. All these marbled crayfish showed the same microsatellite patterns, namely the allele associations 267 bp/271 bp/303 bp at locus PclG-02, 159 bp at PclG-04 and 189 bp/191 bp at PclG-26. The fragment lengths of the alleles of locus PclG-02 overlapped in marbled crayfish (267-303 bp) and *P. fallax* (239-267 bp) but were longer in *P. alleni* (329-384 bp) and shorter in *P. clarkii* (211-228 bp). Marbled crayfish shared two of six alleles with *P. fallax,* namely 267 bp at locus PclG-02 and 159 bp at locus PclG-04, but none with the other species thus confirming the particularly close relationship between *P. fallax* and marbled crayfish.

### 3.3 Ploidy status of marbled crayfish

Martin *et al.* [22] recently used karyological analysis to demonstrate that marbled crayfish has a triploid genome. Our microsatellite analysis confirms this finding. Marbled crayfish generally have the allele association 267 bp/271 bp/303 bp at locus PclG-02 (figure 3*a*), whereas *P. fallax*, *P. alleni* and *P. clarkii* have one or two alleles at this locus, which is consistent with diploid and sexually reproducing species. In an earlier paper, Martin *et al.* [20] have also analysed locus PclG-02 and reported only two alleles of 267 bp and 271 bp. However, a recent re-examination of their material confirmed the presence of the third 303 bp allele (G. Scholtz, personal communication).

**Figure 3.**
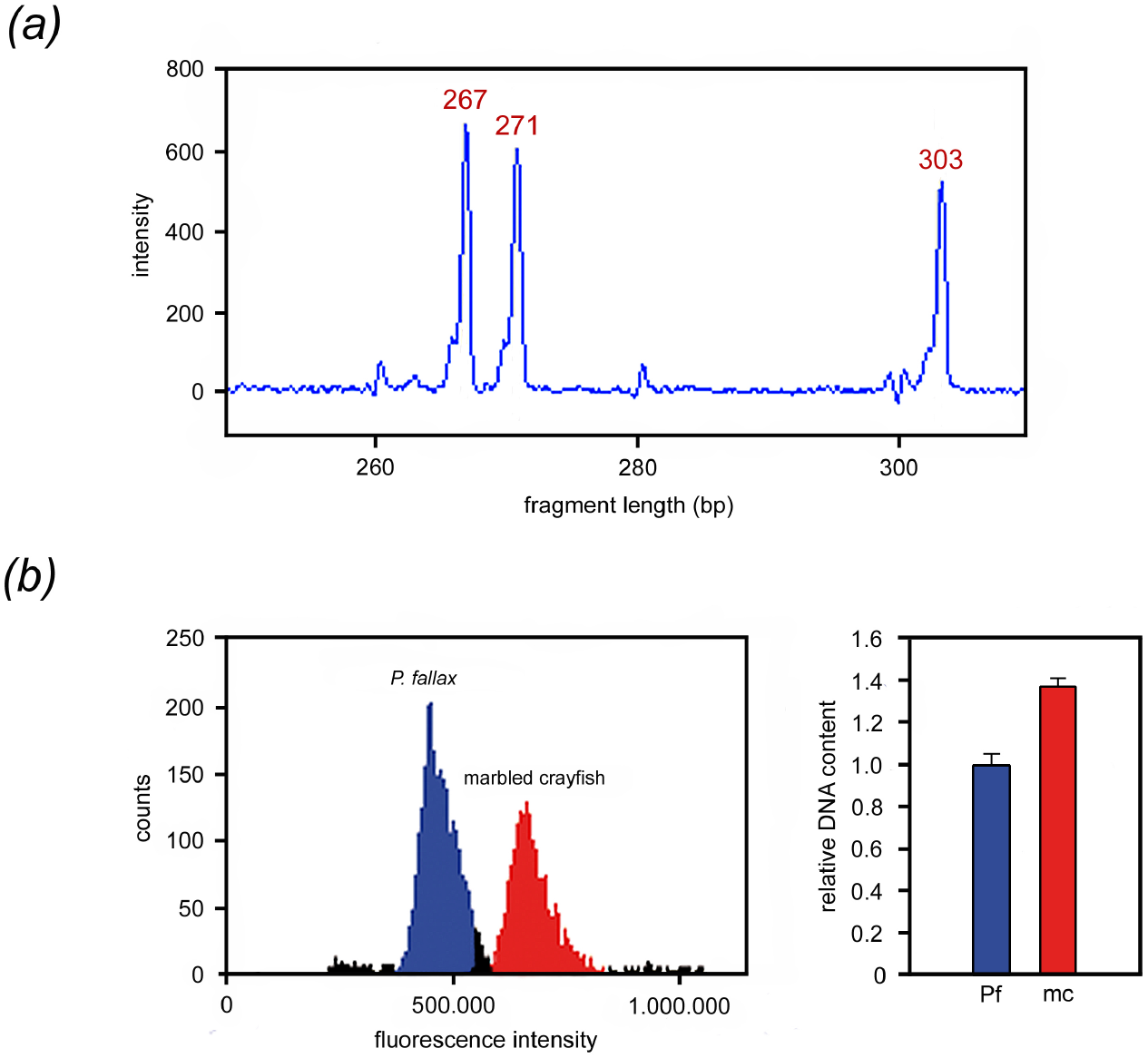
Ploidy status of the marbled crayfish genome. *(a)* Microsatellite locus PclG-02 in marbled crayfish showing a combination of three alleles of 267 bp, 271 bp and 303 bp fragment length. *(b)* Flow cytometry of haemocytes of *P. fallax* (Pf) and marbled crayfish (mc) revealing an approximately 1.4 fold increased DNA content in marbled crayfish. The right panel shows the means and standard deviations of two biological and three technical replicates. Differences are highly significant (p=1.33×10^-7^, Welsh two-sided t-test).

We further corroborated triploidy in marbled crayfish by flow cytometric measurement of the DNA content of haemocytes in marbled crayfish and *P. fallax*. Haemocytes are particularly suitable for this purpose because they are devoid of somatic polyploidization [48]. Our results showed a significant 1.4-fold higher DNA content in the blood cells of marbled crayfish (figure 3*b*), which is consistent with triploidy.

### 3.4 Comparison of DNA methylation between marbled crayfish and *Procambarus fallax*

In order to test if the marbled crayfish and *P. fallax* clusters also differ with respect to epigenetic markers we determined global DNA methylation by mass spectrometry in identically raised and age and size-matched representatives of both crayfish. DNA methylation represents a widely conserved epigenetic mark that is often associated with polyphenism and adaptive phenotypic changes [49,50]. Comparison of three juveniles and selected organs (hepatopancreas, abdominal musculature and ovary) of three adults revealed a consistently and highly significantly reduced level of DNA methylation in marbled crayfish when compared to *P. fallax* (figure 4). The ten *P. fallax* samples together had a DNA methylation level of 2.93±0.15% (mean ± standard deviation) whereas the ten marbled crayfish samples together had a level of only 2.40±0.08%. These results suggest that marbled crayfish have a considerably different DNA methylation pattern.

**Figure 4.**
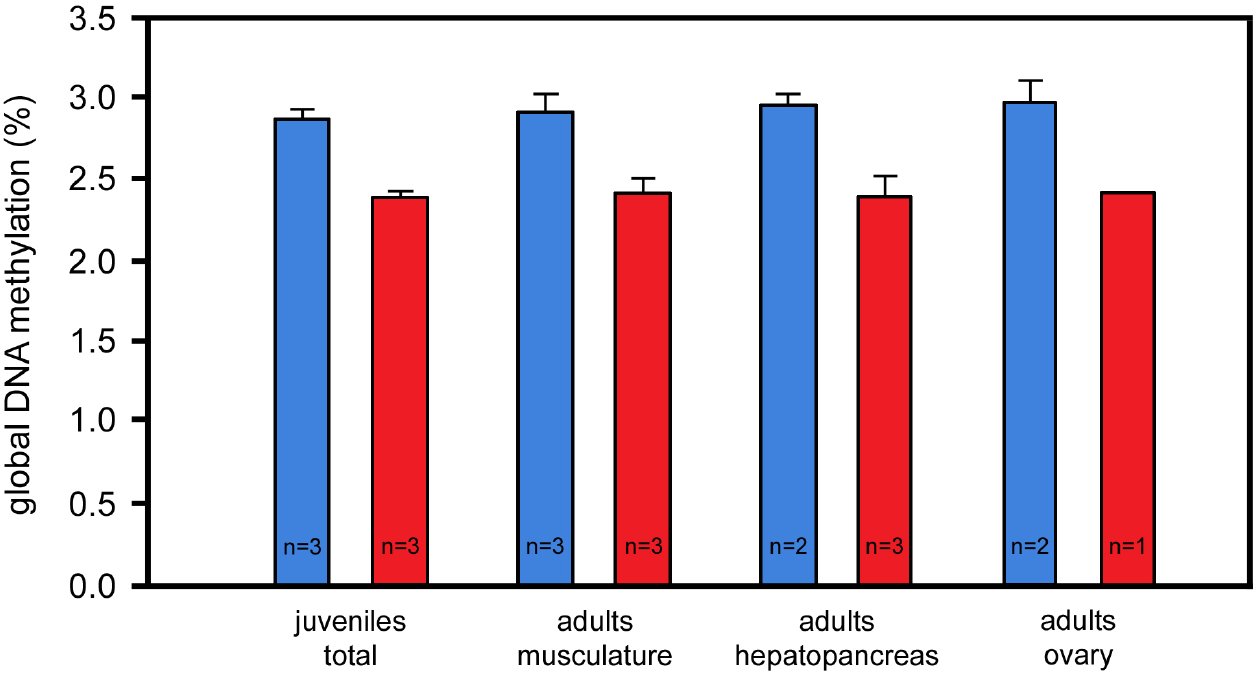
Differences in global DNA methylation between marbled crayfish (red) and *P. fallax* (blue). Analysed were three complete juveniles and major organs of three adult females in each crayfish. Note consistently and significantly greater methylation levels in *P. fallax* (p=1.48×10^-7^ for the sum of all samples, Welsh two-sided t-test). Error bars: standard deviations.

### 3.5 Comparison of morphological characters between marbled crayfish and *P. fallax*

Comparison of the most relevant taxonomic characters of cambarid females [35-37] between marbled crayfish and *P. fallax* corroborated the high degree of morphological similarity between the two crayfish as previously established by Kawai *et al.* [12] and Martin *et al.* [14]. The diagnostically most meaningful trait in females of the genus *Procambarus* is the annulus ventralis, which is bell-shaped with a tilted S-shaped sinus in both marbled crayfish and *P. fallax* (figure 5*a,b*). This typical form is not found in other *Procambarus* species [37] as best exemplified by the differently shaped sperm receptacle of the closely related *P. alleni* (figure 5*c*). The areola, an unpaired structure on the dorsal midline of the carapace, is also very similar in marbled crayfish and *P. fallax* with respect to shape and length-to-width proportion (figure 5*d,e*). The same holds for the cheliped chelae, which closely resemble each other in both crayfish in shape, dentation and setation (figure 5*f,g*), and the coloration pattern, which consists of distinct marmorated spots and dark dorsolateral stripes on carapace and pleon (figure 5*h,i*). Size, form and coloration of the marmoration spots are highly variable not only in the sexually reproducing *P. fallax* but also in the genetically uniform marbled crayfish as a result of stochastic developmental variation [21,51].

**Figure 5.**
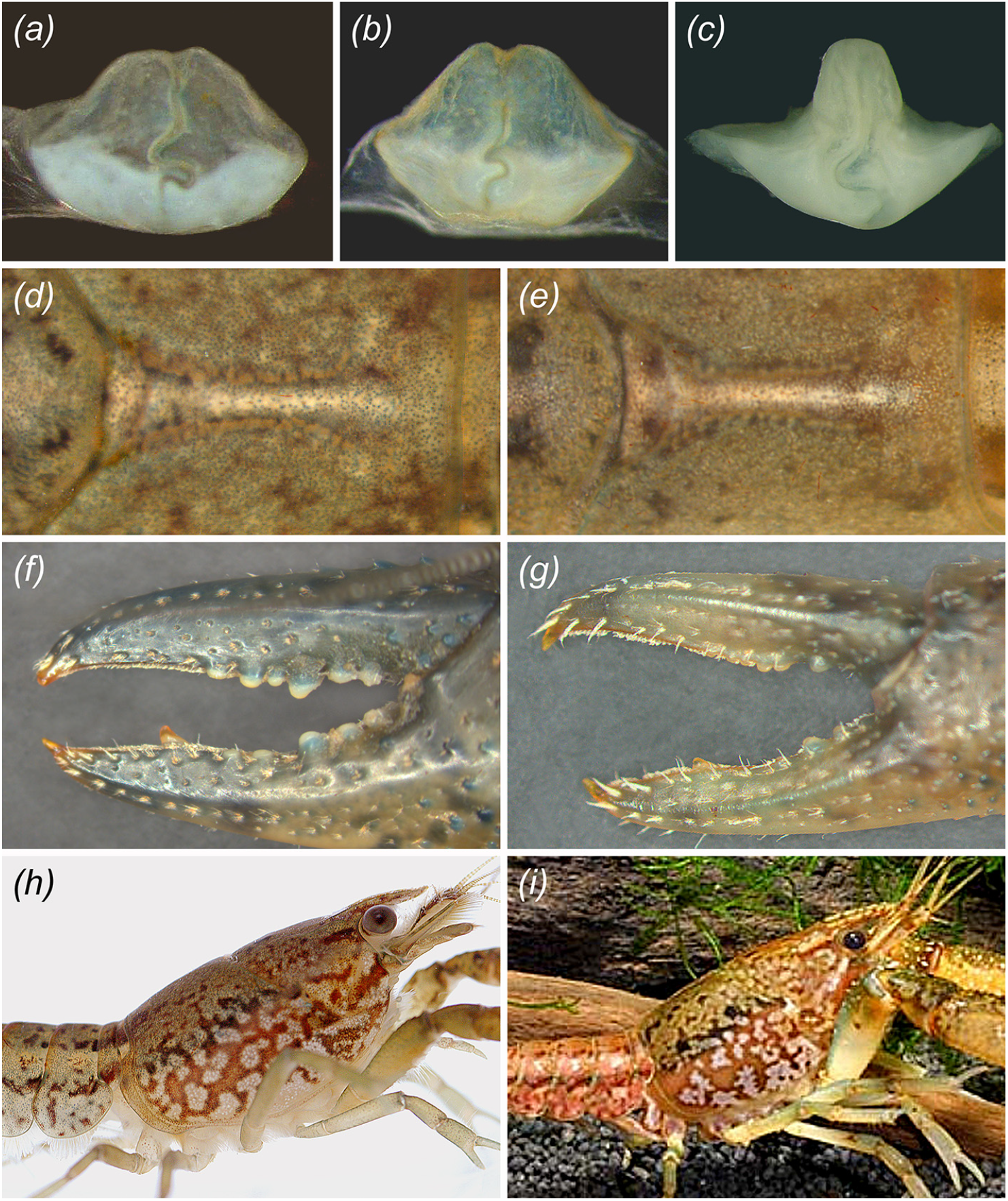
Comparison of morphological characters between marbled crayfish and *P. fallax*. *(a)* Annulus ventralis from exuvia of marbled crayfish. *(b)* Annulus ventralis of *P. fallax*. *(c)* Annulus ventralis of *P. alleni*. Note striking structural difference to sperm receptacles of marbled crayfish and *P. fallax*. *(d)* Areola of marbled crayfish. *(e)* Areola of *P. fallax*. *(f)* Left cheliped of marbled crayfish of 8.4 cm TL. *(g)* Left cheliped of *P. fallax* female of 4.7 cm TL. Form, dentation and setation of the chelae are very similar in both species. *(h)* Coloration of cephalothorax in marbled crayfish. *(i*) Coloration of cephalothorax in *P. fallax* male (photo: C. Lukhaup).

### 3.6 Comparison of life history traits between marbled crayfish and *P. fallax*

In contrast to the morphological characters, life history features like growth and fecundity are markedly different between marbled crayfish and *P. fallax*. Figure 6 gives an example for differences in the speed of growth between identically raised laboratory populations of the same age. At day 250 after hatching, when the first females in both crayfish had reached sexual maturity, mean body weight was almost twice as large in marbled crayfish as in *P. fallax* females.

**Figure 6.**
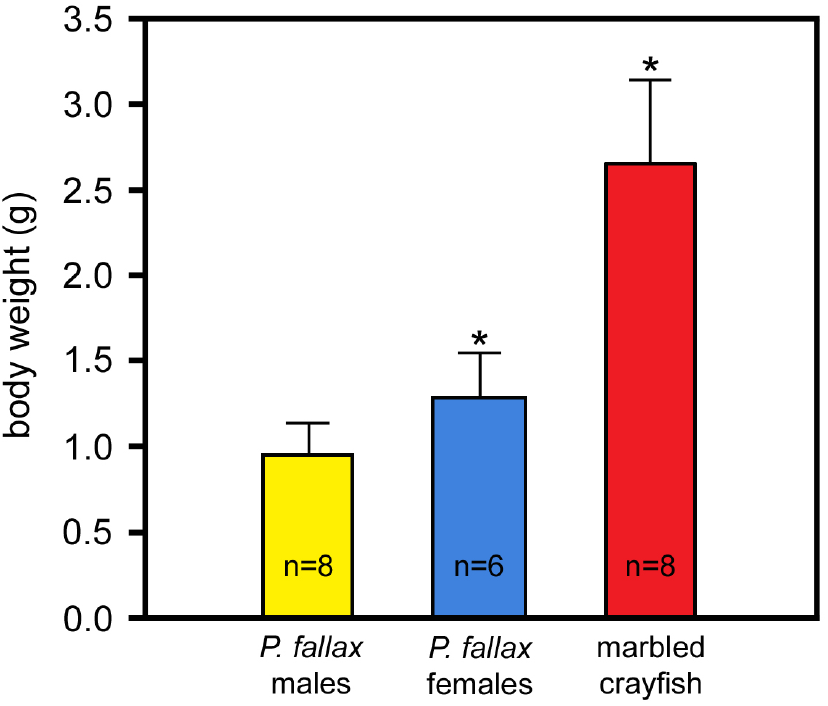
Comparison of growth between marbled crayfish and *P. fallax*. The three groups were reared for 250 days at 20°C under identical conditions and fed with the same food *ad libitum*. The differences between marbled crayfish and *P. fallax* females are highly significant (asterisks; p=2.06×10^-5^; Welsh two-sided t-test). Error bars: standard deviations.

Maximum body and clutch sizes were also markedly higher in marbled crayfish. The largest specimen in our laboratory had a carapace length of 4.9 cm, a total length of 10.3 cm and a body weight of 30.1 g (figure 7*a*). In the wild, the largest of the 1084 marbled crayfish measured [7,12,47, M. Pfeiffer and C. Chucholl, personal communication] was found in Lake Moosweiher and had a CL of 4.9 cm and a weight of 32.0 g [47]. In contrast, the largest of the 4710 wild *P. fallax* examined [36,52-54] had a CL of only 3.4 cm, corresponding to a TL of 7.4 cm and a weight of approximately 11.5 g. The largest clutches of marbled crayfish in the laboratory and the wild consisted of 731 eggs (figure 7*b*) and 724 eggs [47], respectively, which is 5.6 fold higher than the largest clutch of 130 eggs reported for *P. fallax* in literature [53]. The analysis of life history features of the slough crayfish by van der Heiden [54] corroborated that *P. fallax* reaches only rarely a size of more than 6.5 cm TL.

**Figure 7.**
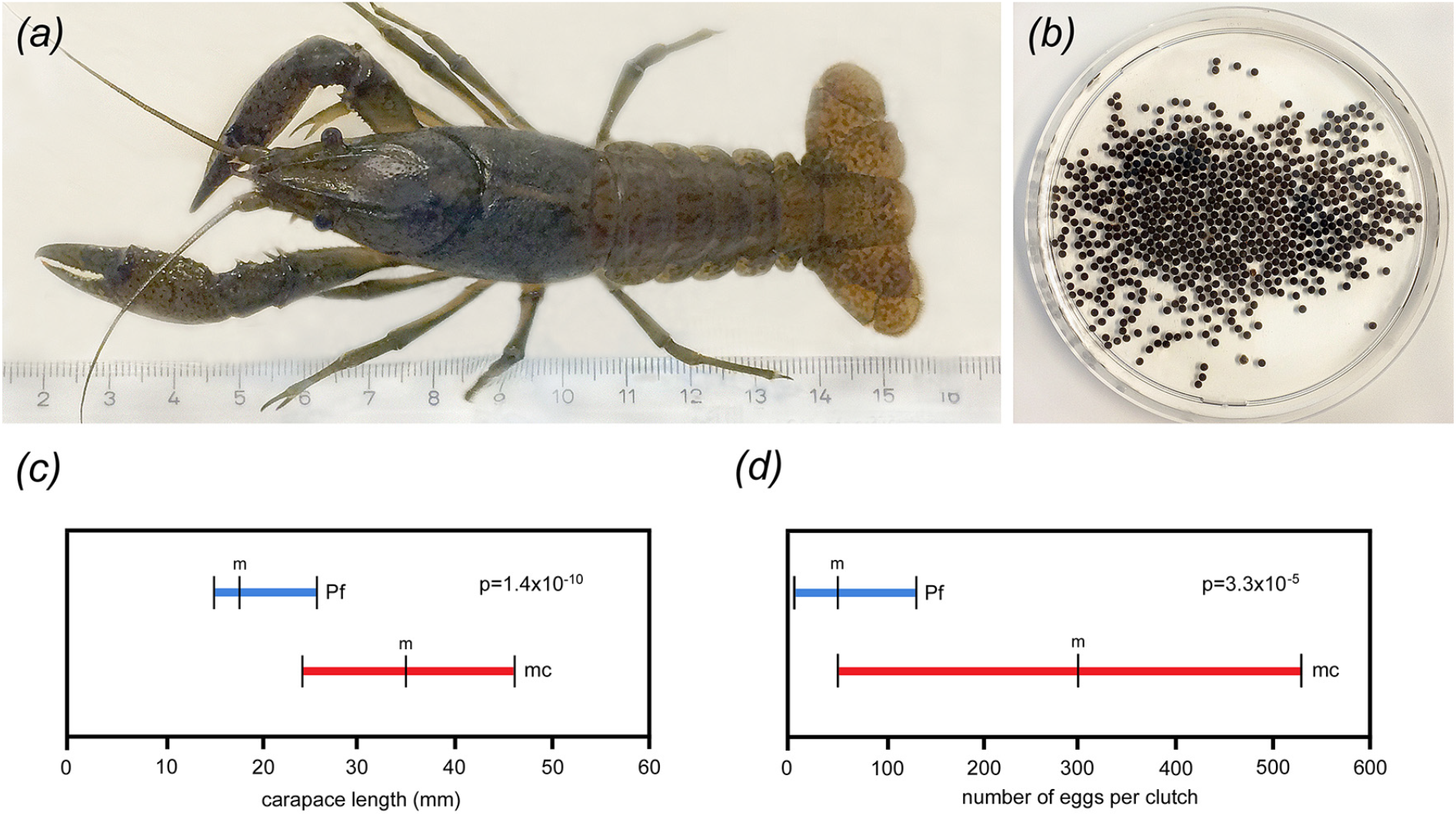
Comparison of body size and fecundity between marbled crayfish and *P. fallax*. *(a)* Largest marbled crayfish from our laboratory having a total length of 10.3 cm. *(b)* Clutch of same specimen consisting of 731 eggs. (*c)* Differences in carapace length between populations of ovigerous marbled crayfish (mc) and *P. fallax* females (PF) from comparable climatic regions. Data for marbled crayfish (n=57) was obtained in Madagascar [7] and data for *P. fallax* (n=27) was obtained in Florida [53]. Horizontal bars indicate ranges and vertical lines indicate mean values (m) and lower and upper range limits. The difference between marbled crayfish and *P. fallax* females is highly significant as indicated by the p-value. (*d*) Differences in clutch size between the same populations as in (*c*). The difference is highly significant as indicated by the p-value. For statistical calculations, the standard deviation was taken as half the range, and a Bonferroni adjustment for multiplicity was applied.

The differences in growth and fecundity between marbled crayfish and *P. fallax* were also confirmed by the analysis of published data for egg-carrying females from comparable climatic regions. Ovigerous marbled crayfish from Madagascar had a mean CL of 3.5 cm, a mean TL of 7.4 cm and a mean clutch size of 300 eggs [7], whereas ovigerous *P. fallax* from the Everglades National Park in Florida had a mean CL of 1.8 cm, a mean TL of 3.8 cm and a mean clutch size of 41 eggs only [53], indicating that body size and fecundity is significantly increased in marbled crayfish (figure 7*c*,*d*). These findings identify important phenotypic differences between marbled crayfish and *P. fallax* that have not been recognized previously.

## 4. Discussion

Our results demonstrate that marbled crayfish meets all criteria for asexual speciation [25-28]. It is separated from the mother species, *P. fallax*, by reproductive isolation, significant genomic and epigenetic differences and superior life history traits. Our data further support a single origin. In addition, all populations known to date live outside the natural range of *P. fallax*, suggesting geographical isolation. They are unified in one cluster by common phenotypic, genetic and epigenetic characteristics, despite their broad geographical distribution. These commonalities and differences towards *P. fallax* make it very likely that the marbled crayfish and slough crayfish clusters will evolve differently, which is the main criterion for erecting an asexual species [26]. Martin *et al.* [14] have previously suggested that marbled crayfish should be considered as an independent species when a single origin and/or regional populations in the wild have been established. Our findings clarify the former issue and provide additional evidence for cytogenetic, genetic and phenotypic differences between marbled crayfish and *P. fallax*. As such, marbled crayfish should now be named *Procambarus virginalis*, as suggested previously [14]. The formal description of marbled crayfish as a new species will be detailed in a separate publication.

Marbled crayfish appeared first in 1995 in the German aquarium trade. Thereafter, aquarists have propagated it in captivity, and since about 2003, releases have resulted in the establishment of thriving wild populations in Central Europe and Madagascar [5,7-9,12,47]. The “mega-population” [46] in innumerable aquarium tanks on various continents and the known wild populations are apparently all descendants of the single clone or single individual that was introduced in Germany in 1995. Our results confirm this single origin by the identity of the mitochondrial genomes and microsatellite patterns in samples of captive and wild populations. One of the samples analysed in our study, the Heidelberg specimen, can be directly traced back to the year 1995 and to the oldest marbled crayfish for which written records exist (F. Steuerwald, personal communication).

It is unknown whether marbled crayfish emerged in the natural range of *P. fallax* or in captivity. Scholtz [4], Faulkes [5] and Martin [46] summarized possible scenarios for the first alternative including hybridization with coexisting *Procambarus* species and geographic parthenogenesis. These authors and Chucholl [9] also stressed that in captivity there were many more candidates for hybridization than the naturally coexisting six *Procambarus* species [36,52] because crayfish were popular pets already in the 1990s. Faulkes [5] emphasized that all surveys on *P. fallax* in Florida and Southern Georgia revealed males and females arguing against the presence of pure marbled crayfish populations in the natural range of *P. fallax*. Moreover, none of the articles on wild *P. fallax* [36,45,52-54] mentioned specimens above 7.4 cm TL, which would again support the absence of primary populations of marbled crayfish. In sympatric populations, small and medium-sized marbled crayfish and *P. fallax* females would be indistinguishable by morphological criteria alone. However, by the use of genetic markers marbled crayfish could now be identified. Particularly useful is the highly specific tri-allelic microsatellite locus PclG-02, which could be assayed in large samples with reasonable expenditure. However, time for the detection of primary populations may be limited because marbled crayfish are already widespread in American aquaria [11] and their release into the natural range of *P. fallax* would render the search for primary populations of marbled crayfish impossible.

Our crossbreeding experiments with marbled crayfish, *P. fallax* and *P. alleni* revealed that marbled crayfish and *P. fallax* still recognize each other as sexual partners but not marbled crayfish and *P. alleni*. Recognition of sexual partners in crayfish is mainly based on chemical signatures of the urine but may also include visual and tactile cues [38,39]. Marbled crayfish and *P. fallax* copulate readily with each other. However, the progeny of such pairings are pure marbled crayfish resulting from parthenogenesis. These findings demonstrate reproductive isolation and suggest that the reproductive barrier is set at the cytogenetic rather than the behavioural level. Mechanical barriers can be largely excluded because the sperm receptacles are structurally very similar in marbled crayfish and *P. fallax* females and because we have repeatedly observed insertion of the male gonopods into the annulus ventralis of marbled crayfish. We attempted to directly prove sperm transfer by analysing moulted sperm receptacles of females that had successfully produced offspring. However, we did not find any sperm remnants neither in marbled crayfish nor *P. fallax* females.

The morphological features and microsatellite patterns strongly suggest that marbled crayfish originated by autopolyploidization and not by hybridization with a closely related species, which is by far the most frequent cause of triploidy in animals [55-58]. Typically, hybrids between two crayfish species are clearly recognizable because of their intermediate morphological characters [59,60]. However, marbled crayfish do not show such hybrid features [12,14, this study]. Conversely, autopolyploids are usually morphologically similar to their diploid progenitors [61], and the morphological similarity between marbled crayfish and *P. fallax* is therefore consistent with autopolyploidization. There is also no evidence for hybridization on the genetic level and no strong bias towards heterozygosity in the microsatellite pattern, which would be typical for hybrids [62,63]. Of the seven microsatellite loci that were investigated in marbled crayfish so far, three were homozygous and four were heterozygous [20,21, this study], thus largely excluding allopolyploidization for marbled crayfish. Furthermore, Martin and colleagues have recently shown that the nuclear elongation factor 2 (EF-2) gene is identical in marbled crayfish and *P. fallax* but differs from other *Procambarus* species like *P. alleni*, *P. clarkii*, *P. acutus* and *P. liberorum* [22]. These findings provide additional support for the origin of marbled crayfish by autopolyploidization.

We admit that the presence of three alleles, as observed in locus PclG-02 in marbled crayfish, can be interpreted to reflect an origin by hybridization. However, such a pattern can also occur in autopolyploids, namely when an unreduced diploid egg is fertilized by a sperm from the same species, or alternatively, by simultaneous fertilization of a haploid egg by two sperms with different alleles. In shrimp, fish and bivalve aquaculture, autopolyploid triploids with tri-allelic loci are artificially produced by the prevention of polar body I extrusion in fertilized eggs either by temperature shock or chemicals like 6-dimethylaminopurine [64,65]. Marbled crayfish may thus have arisen by a heat or cold shock in the sensitive phase of egg development in a captive *P. fallax* female, possibly during transportation.

The origin of parthenogenesis in marbled crayfish is probably a by-product of polyploidization but the causal relationship of polyploidy and parthenogenesis is not yet understood [46]. Infectious parthenogenesis by the feminizing bacterium *Wolbachia*, which is widespread in crustaceans [66], was excluded by the use of molecular probes for the parasite [2]. In plants, it was shown that polyploidy per se can have an immediate impact on the reproductive biology of a species [67]. In animals, however, obligate parthenogenesis is relatively rare. It has been described in some asexual invertebrate families and a few vertebrate hybrids [26,68-71] and is mostly associated with allopolyploidy. Autopolyploidy is much less common and is usually not associated with parthenogenesis, perhaps with the exception of some high arctic ostracods and polyploid populations of the brine shrimp *Artemia parthenogenetica* [72,73]. Artificially induced autopolyploid shrimp and fish are usually sterile [74], making the combination of autopolyploidy and parthenogenesis in marbled crayfish rather unique.

Polyploids often have life history traits that are different from those of the parent species. Growth, number of offspring and other quantitative traits can either be decreased or increased when compared to the diploid ancestors [75-77]. In marbled crayfish, growth, maximum body size and fecundity were significantly increased when compared to *P. fallax*, whereas the time of sexual maturity was similar (7,36,47,54, this study). Longevity may also be increased in marbled crayfish. Maximum age so far recorded is 1610 days in marbled crayfish [19] and 980 days in *P. fallax* (Z. Faulkes, personal communication). These superior fitness traits, together with parthenogenetic reproduction, are probably causative for the remarkable success of marbled crayfish as an invasive species in Central Europe and Madagascar [7-9,47]. Chucholl [9] calculated an almost double FI-ISK (Freshwater Invertebrate Invasiveness Scoring Kit) score for marbled crayfish when compared to *P. fallax*, making it a high risk species for Central Europe. Moreover, Feria and Faulkes [78] predicted with climate and habitat based Species Distribution Models that marbled crayfish could inhabit a larger geographical area than its mother species *P. fallax* when released in the southern states of the USA, thus illustrating the ecological superiority of marbled crayfish.

In allopolyploids, the increase of life history traits is usually explained as the result of heterozygosity, which is well known as heterosis effect or hybrid vigor [79,80]. However, this explanation is not applicable for autopolyploids because autopolyploidization enhances only the copy number of already existing genes. However, novel traits do not necessarily require new genes or new developmental pathways to come into being but can instead arise from recruitment of already existing developmental processes into new contexts [81,82]. Thus, trait alteration in marbled crayfish may have been caused by altered gene dosage, the rearrangement of gene-networks and the modulation of gene expression by changes in epigenetic regulation.

Changes in epigenetic regulation can be deduced from the significantly reduced level of global DNA methylation in marbled crayfish when compared to *P. fallax*. DNA methylation is an epigenetic mechanism that considerably affects plant and animal phenotypes [49,50,83]. It is responsive to environmental and genomic stresses including polyploidization [50] and might thus contribute to speciation in polyploids. In plants, the increase or reduction of global DNA methylation after autopolyploidization is well known [61,84]. It is also well established that DNA methylation and other epigenetic mechanisms contribute to the establishment of reproductive barriers [85,86] and the expression of hybrid vigor in allopolyploid plants [87]. In marbled crayfish, epigenetic mechanisms may thus have been involved in the acquisition of novel fitness traits.

Chen *et al.* [88] reported that polyploidization is often accompanied or followed by intense rearrangements in the genome, which stabilize the new lineage. These rearrangements, which are associated with epigenetic changes, can include loss of DNA. For example, in synthetic autopolyploids of annual phlox, *Phlox drummondii*, an immediate loss of 17% of total DNA has been observed with a further reduction of up to 25% upon the third generation [89]. Such mechanisms may also have operated during transition from *P. fallax* to marbled crayfish and might explain why triploid marbled crayfish have only a 1.4-fold rather than a 1.5-fold increased DNA content when compared with its diploid mother species.

Speciation by autopolyploidization is a special case of chromosomal speciation that is well-known in plants [61] but virtually unknown in animals. Chromosomal speciation is a complementary concept to the better known speciation by changes in allele frequency distribution and can result in the almost instantaneous production of new species and phenotypic novelty within one generation [90-92]. This “saltational speciation” or “saltational evolution” [93-95] has largely been ignored by gradualism-based Modern Synthesis, which may be due to its rarity in animals, the lack of mechanistic understanding and the dearth of suitable models. Marbled crayfish represents a contemporary animal example of autopolyploid speciation, which likely started about 20-30 generations ago. Comparative genome and epigenome sequencing approaches will be required to fully understand the genetic and epigenetic differences between both species.

## 5. Conclusion

Marbled crayfish can be regarded as a new species that originated from *P. fallax* by triploidization and concomitant epigenetic alterations, as shown by our combined morphological, behavioural, genetic and epigenetic analysis. Marbled crayfish is morphologically very similar to its mother species but has superior fitness traits. Genetic data suggest an instantaneous speciation by autopolyploidization and parallel change of the mode of reproduction from gonochorism to parthenogenesis. The young evolutionary age of marbled crayfish, which is possibly three decades or less, may offer the possibility to identify key events for this type of speciation. The combination of autopolyploidy and obligate parthenogenesis is common in plants but very rare in animals. Thus, the *P. fallax*-marbled crayfish pair provides an interesting new model system to study asexual speciation and saltational evolution in animals and to determine how much genetic and epigenetic change is necessary to create a new species.

## Acknowledgement

We thank Michael Pfeiffer (Gobio, March-Hugstetten, Germany) and Christoph Chucholl (Fisheries Research Station Baden-Württemberg, Langenargen, Germany) for providing marbled crayfish from Lake Moosweiher and for information on the biology of marbled crayfish in this lake, Frank Glaw (Zoologische Staatssammlung, Munich, Germany) and Miguel Vences (Braunschweig University of Technology, Germany) for the Madagascar sample, the Bundesamt für Umwelt (Bern, Switzerland) for the *Procambarus clarkii* samples, Frank Steuerwald (KABS, Waldsee, Germany) for information on the oldest known marbled crayfish, Chris Lukhaup (Hinterweidenthal, Germany) for figure 5*i*, Thomas Carell (Ludwig-Maximilians-University, Munich, Germany) for providing [D_3_]-dm^5^C internal standard for mass spectrometry, Günter Raddatz and Carine Legrand (DKFZ) for statistical help, the DKFZ Flow Cytometry and Genomics and Proteomics Core Facilities for flow cytometry and DNA sequencing services, and Gerhard Scholtz (Humboldt University, Berlin, Germany), Bronwyn W. Williams (North Carolina Museum of Natural Sciences, Raleigh, USA) and Zen Faulkes (University of Texas-Pan American, Edinburg, USA) for valuable comments that improved the manuscript.

## Authors’ contributions

G.V. conceived of the study, participated in the design of the study, sampled the tissues, performed the cross-breeding experiments and analysed the morphological and life history data; C.F. carried out the assembly and analysis of mitochondrial genome sequences and the determination of DNA contents by flow cytometry; K.H. maintained laboratory crayfish cultures and prepared DNA samples; A.S., J.P. and R.S. performed the analysis of the microsatellite markers; K.S and M.H. carried out the mass spectrometric measurement of DNA methylation; F.L. participated in the design of the study and coordinated the study. G.V. and F.L. wrote the manuscript. All authors revised the manuscript and gave final approval for publication.

## Data accessibility

The mitochondrial DNA sequences have been deposited in GenBank under the accession numbers KT074363, KT074364 and KT074365.

## Ethics statement

All crayfish experiments were performed by approval of the institutional animal welfare committee, in compliance with local standards and guidelines.

